## Supplementary material for "Collective search with finite perception: transient dynamics and search efficiency"

(Dated: November 13, 2018)

### S1. COMPUTATION OF THE EFFECTIVE GRADIENT UNDER LIMITED SPATIAL EXPLORATION

Here, we compute the effective gradient of a searcher, while relaxing the assumption that the searchers have explored the whole domain. In this case, the effective gradient of a searcher starting at $x_0$, will be given by the average gradient in the explored domain up to the search time $T$. In one dimension, this is

$$\begin{aligned}\langle \nabla S(x)\rangle_T &= \frac{1}{x_{\mathrm{ms}}(T)}\int_{x_0}^{x_0+x_{\mathrm{ms}}(T)} \frac{S_1\left(x+x_{\mathrm{ms}}(\tau)\right)-S_1(x)}{x_{\mathrm{ms}}(\tau)}dx \\ &= \frac{A}{x_{\mathrm{ms}}(T)x_{\mathrm{ms}}(\tau)}\int_{x_0}^{x_0+x_{\mathrm{ms}}(T)} \exp\left(-\frac{x^2}{2\sigma^2}\right)\left[\exp\left(-\frac{2xx_{\mathrm{ms}}(\tau)+x_{\mathrm{ms}}(\tau)^2}{2\sigma^2}\right)-1\right]dx, \qquad \text{(S1)}\end{aligned}$$

where $A = 1/\sqrt{2\pi\sigma^2}$ is a normalisation constant.

Then, we use the assumption $x_{\mathrm{ms}}(\tau) \ll x_{\mathrm{ms}}(T)$, and evaluate (S1) asymptotically to first order in two extreme cases: when $x_{\mathrm{ms}}(T) \ll \sigma$ and $\sigma \ll x_{\mathrm{ms}}(T)$. First, for limited search times we have that $x_{\mathrm{ms}}(\tau) \ll x_{\mathrm{ms}}(T) \ll \sigma \ll 1$. Therefore, we may write

$$\begin{aligned}\langle \nabla S(x)\rangle_T &= \frac{A}{x_{\mathrm{ms}}(T)}\int_{x_0}^{x_0+x_{\mathrm{ms}}(T)} \exp\left(-\frac{x^2}{2\sigma^2}\right)\left[-\frac{x}{\sigma^2}+\mathcal{O}\left(\frac{x_{\mathrm{ms}}(\tau)}{\sigma^2}\right)\right]dx \\ &= \frac{A}{x_{\mathrm{ms}}(T)}\int_{x_0}^{x_0+x_{\mathrm{ms}}(T)} \exp\left(-\frac{x^2}{2\sigma^2}\right)\left[-\frac{x}{\sigma^2}\right]dx + \mathcal{O}\left(\frac{x_{\mathrm{ms}}(\tau)}{\sigma^2}\right) \\ &= \frac{A}{x_{\mathrm{ms}}(T)}\left[\exp\left(-\frac{(x_0+x_{\mathrm{ms}}(T))^2}{2\sigma^2}\right)-\exp\left(-\frac{x_0^2}{2\sigma^2}\right)\right]+\mathcal{O}\left(\frac{x_{\mathrm{ms}}(\tau)}{\sigma^2}\right). \qquad \text{(S2)}\end{aligned}$$

Therefore, when $x_0 \sim 1$, we obtain that $\langle \nabla S(x)\rangle_T = \mathcal{O}\left(x_{\mathrm{ms}}(\tau)/\sigma^2\right)$, meaning that for cells far from the origin the contribution to motion by gradient-driven advection can be neglected on short time scales.

Meanwhile, close to the origin, $x_0 \approx 0$, we may expand the argument in the exponential in (S2), namely

$$\begin{aligned}\langle \nabla S(x) \rangle_T &= \frac{A}{x_{\mathrm{ms}}(T)} \left[ -\frac{(x_0 + x_{\mathrm{ms}}(T))^2}{2\sigma^2} + \frac{x_0^2}{2\sigma^2} \right] + \mathcal{O}\left( \frac{x_{\mathrm{ms}}(\tau)}{\sigma^2} \right) \\ &= \frac{A x_{\mathrm{ms}}(T)}{\sigma^2} + \mathcal{O}\left( \frac{x_{\mathrm{ms}}(\tau)}{\sigma^2} \right) + \mathcal{O}\left( \frac{x_0}{\sigma^2} \right) \\ &\propto \frac{1}{\sigma^3} + \mathcal{O}\left( \frac{x_{\mathrm{ms}}(\tau)}{\sigma^2} \right) + \mathcal{O}\left( \frac{x_0}{\sigma^2} \right). \end{aligned} \tag{S3}$$

Second, for large search times, the search behaviour is independent of the starting point thus we may set $x_0 = 0$ in (S1). If in addition, we have that $x_{\mathrm{ms}}(\tau) \ll \sigma \ll x_{\mathrm{ms}}(T) \sim 1$, then the first order asymptotic solution to (S1) reads:

$$\begin{aligned}\langle \nabla S(x) \rangle_T &= \frac{A}{x_{\mathrm{ms}}(T)} \int_0^{x_{\mathrm{ms}}(T)} \exp\left( -\frac{x^2}{2\sigma^2} \right) \left[ -\frac{x}{\sigma^2} \right] dx + \mathcal{O}\left( \frac{x_{\mathrm{ms}}(\tau)}{\sigma^2} \right) \\ &= -\frac{A}{x_{\mathrm{ms}}(T)} \left[ \exp\left( -\frac{x_{\mathrm{ms}}^2(T)}{2\sigma^2} \right) - 1 \right] + \mathcal{O}\left( \frac{x_{\mathrm{ms}}(\tau)}{\sigma^2} \right) \\ &\propto \frac{1}{\sigma} + \mathcal{O}\left( \frac{x_{\mathrm{ms}}(\tau)}{\sigma^2} \right) + \mathcal{O}\left( \frac{\sigma}{x_{\mathrm{ms}}(T)} \right). \end{aligned} \tag{S4}$$

If on the other hand $\sigma \ll x_{\mathrm{ms}}(\tau) \ll x_{\mathrm{ms}}(T) \sim 1$, then we get that

$$\begin{aligned}\langle \nabla S(x) \rangle_T &= 2A \left[ \left( 1 - e^{-x_{\mathrm{ms}}(T)/(2\sigma^2)} \right) - \frac{x_{\mathrm{ms}}(\tau)}{4\sigma} e^{-x_{\mathrm{ms}}(T)/(2\sigma^2)} \right] + \mathcal{O}(x_{\mathrm{ms}}^2(\tau)) \\ &\propto \frac{1}{\sigma} + \mathcal{O}\left( \frac{x_{\mathrm{ms}}^2(\tau)}{\sigma^2} \right) + \mathcal{O}\left( \frac{\sigma}{x_{\mathrm{ms}}(T)} \right). \end{aligned} \tag{S5}$$

Thus, to first order, when $x_{\mathrm{ms}}(T) \ll \sigma$ we have that $\langle \nabla S(x) \rangle_T \propto \sigma^{-3}$, while for $\sigma \ll x_{\mathrm{ms}}(T)$ we have that $\langle \nabla S(x) \rangle_T \propto \sigma^{-1}$.

### S2. GRADIENT-SENSING IS THE OPTIMAL STRATEGY IN LINEAR STIMULANT LANDSCAPES

In linear stimulant landscapes the local gradient already contains complete information about the whole landscape. Therefore, the intuition is that in this case, the solution of the optimal foraging model will be invariant under $\tau$, i.e. $\rho_{\mathrm{ON}}(x,t;\tau)=\rho_{\mathrm{ON}}(x,t)$.

To formalise this, we solve the ON model (Eq. (11) in the paper) for $S_1=\alpha x$. As derived in Appendix B (Eqs. (B3a) and (B3b) in the paper), the optimality conditions of (11) read

$$\partial_s m+\nabla\cdot(mu)=0,\ \ m(\cdot,0)=\rho(\cdot,k\tau) \tag{B3a}$$

$$\partial_s\phi+u\cdot\nabla\phi+\frac{|u|^2}{2}=0,\ \ \phi(\cdot,\tau)=-\frac{\delta\mathcal{F}[m]}{\delta m}\bigg|_{s=\tau}, \tag{B3b}$$

with $u=-\nabla\phi$ and $\mathcal{F}[m]=\int_\Omega\big(m\log m-\mathrm{Pe}_1 mS_1-\frac{\mathrm{Pe}_2}{2}m(\log|m|*m)\big)\,dx$. Note that since we are interested in the effect of the landscape $S_1$ on the drift velocity, we let $\mathrm{Pe}_2=0$ (i.e. the searchers are non-interacting) and $\int_\Omega\rho\log\rho dx=0$. Considering the HJ equation (B3b) we can then write

$$\partial_s\phi-\frac{1}{2}|\nabla\phi|^2=0,\qquad \phi(x,\tau)=-\mathrm{Pe}_1S_1(x), \tag{S6}$$

Equation (S6) can be solved by the general method of characteristics ([1], ch. 3), as follows. Define the parametrisation of space ($x$) and time ($s$) in terms of $\mu$ and $\nu$, respectively, and let

$$\begin{aligned}p(\mu,\nu)&:=\partial_s\phi(x(\mu,\nu),s(\mu,\nu))\\ q(\mu,\nu)&:=\nabla\phi(x(\mu,\nu),s(\mu,\nu))\\ z(\mu,\nu)&:=\phi(x(\mu,\nu),s(\mu,\nu)).\end{aligned}$$

Then we may write (S6) as:

$$\begin{cases}F(x,s,z,p,q)=p-\frac{|q|^2}{2}=0\\ z(x,\tau)=-\mathrm{Pe}_1S_1(x)\end{cases} \tag{S7}$$

The characteristic equations and the corresponding boundary conditions are:

$$\frac{dx}{d\nu} = \frac{\partial F}{\partial q} = -q; \qquad x(\mu,0) = \mu \tag{S8a}$$

$$\frac{ds}{d\nu} = \frac{\partial F}{\partial p} = 1; \qquad s(\mu,0) = \tau \tag{S8b}$$

$$\frac{dz}{d\nu} = p\frac{\partial F}{\partial p} + q\frac{\partial F}{\partial q} = p - q^2; \qquad z(\mu,0) = -\mathrm{Pe}_1 S_1(\mu) \tag{S8c}$$

$$\frac{dp}{d\nu} = -\frac{\partial F}{\partial s} - p\frac{\partial F}{\partial z} = 0; \qquad p(\mu,0) = \psi_1(\mu) \tag{S8d}$$

$$\frac{dq}{d\nu} = -\frac{\partial F}{\partial x} - q\frac{\partial F}{\partial z} = 0; \qquad q(\mu,0) = \psi_2(\mu). \tag{S8e}$$

Here $\psi_1$ and $\psi_2$ are boundary conditions for $p$ and $q$, respectively, which must therefore satisfy (S7)

$$F(x, s, z, \psi_1, \psi_2) = 0,$$

along with the the compatibility condition

$$\frac{d}{d\mu} z(x(\mu,\nu), s(\mu,\nu))\Big|_{\nu=0} = \nabla z \frac{\partial}{\partial \mu} x(\mu,\nu)\Big|_{\nu=0} + \frac{\partial z}{\partial s}\frac{\partial}{\partial \mu} s(\mu,\nu)\Big|_{\nu=0}.$$

Hence the following two conditions must hold

$$\begin{cases} \frac{d}{d\mu} z(\mu,0) = q\frac{\partial}{\partial \mu} x(\mu,0) + p\frac{\partial}{\partial \nu} s(\mu,0) \\ \psi_1 - \frac{|\psi_2|^2}{2} = 0. \end{cases}$$

Using the boundary conditions in (S8), we obtain that $\psi_1 = |\partial_\mu S_1(\mu)|^2/2$ and $\psi_2 = -\partial_\mu S_1(\mu)$. Then, integrating the characteristic equations (S8) we obtain

$$x = \partial_\mu S_1(\mu)\nu + \mu \tag{S9}$$

$$s = \tau - \nu \tag{S10}$$

$$z = \frac{|\partial_\mu S_1(\mu)|^2}{2}\nu - \mathrm{Pe}_1 S_1(\mu), \tag{S11}$$

which need to be inverted to express $\mu, \nu, z$ in terms of $x, s$.

*Particularisation for linear stimulant landscapes* — For linear landscapes $S_1(\mu) = \alpha\mu$, it is possible to invert the above equations to obtain $\phi = z(x,s) = -\alpha \mathrm{Pe}_1 x + f(s)$. Using the optimality condition, we find that the mean chemotactic velocity is $u(x,s) = -\nabla\phi = \alpha\mathrm{Pe}_1$, independent of $\tau$, as we set out to show.

In conclusion, under linear stimulant landscape local gradient information characterises the field globally and there is no benefit from having a finite spatial horizon.

### S3. SUPPLEMENTARY FIGURES

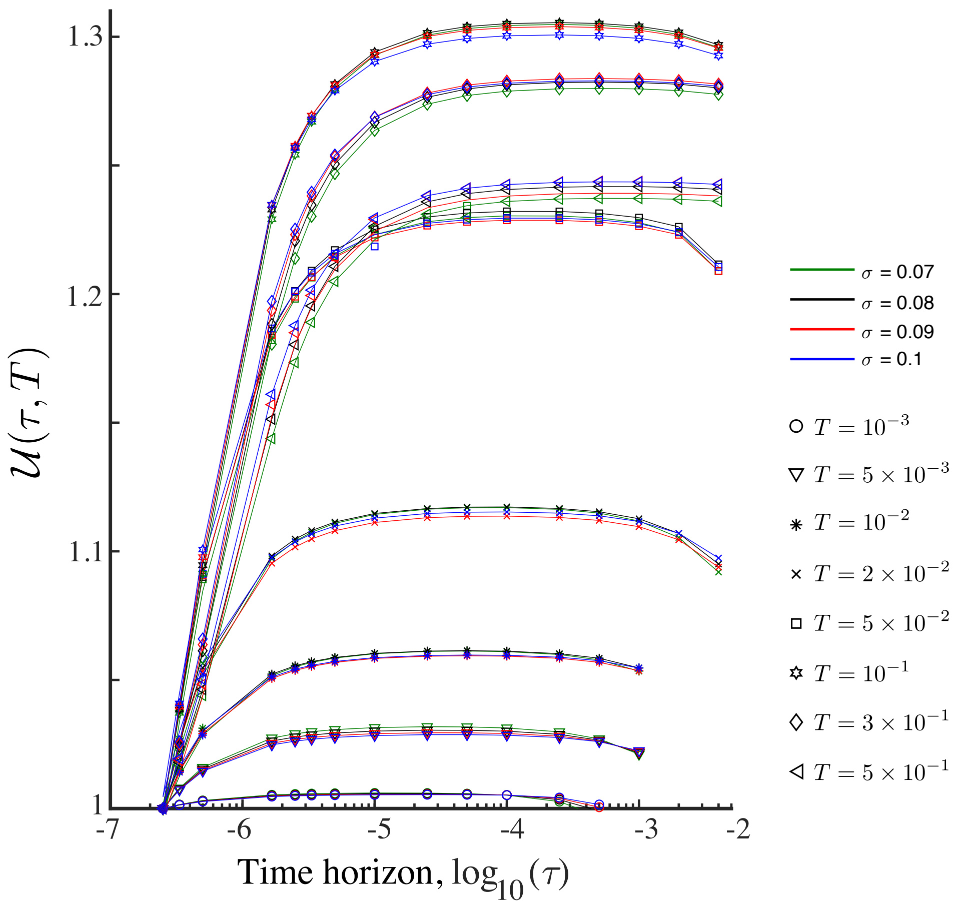


Figure S1: Relative population fitness against time horizon at fixed search times $T$ and different environmental length scales $\sigma$ (see legend). Markers are obtained by renormalising the Péclet number as in (23), simulating a population evolution in the given stimulant landscape using the ON model (Eq. (11) in the paper) and computing $\mathcal{U}(\tau, T)$ from (17). The maxima of the curves $\mathcal{U}^*(T)$ are used to compute Fig. 4d in the paper.

### REFERENCES


[1] L. C. Evans, *Partial Differential Equations*, edited by F. Brezzi, P. Colli Franzone, U. Gianazza, and G. Gilardi, Vol. 4 (AMS, 2010).
